# Neutrophils encompass a regulatory subset suppressing T cells in apparently healthy cattle and mice

**DOI:** 10.1101/2020.11.02.364885

**Authors:** Marion Rambault, Émilie Doz-Deblauwe, Yves Le Vern, Florence Carreras, Patricia Cunha, Pierre Germon, Pascal Rainard, Nathalie Winter, Aude Remot

## Abstract

Neutrophils that reside in the bone marrow are switly recruited from circulating blood to fight infections. For a long time, these first line defenders were considered as microbe killers. However their role is far more complex as cross talk with T cells or dendritic cells have been described for human or mouse neutrophils. In cattle, these new roles are not documented yet. We identified a new subset of regulatory neutrophils that is present in the mouse bone marrow or circulate in cattle blood under steady state conditions. These regulatory neutrophils that display MHC-II on the surface are morphologically indistinguishable from classical MHC-II^neg^ neutrophils. However MHC-II^pos^ and MHC-II^neg^ neutrophils display distinct transcriptomic profiles. While MHC-II^neg^ and MHC-II^pos^ neutrophils display similar bacterial phagocytosis or killing activity, MHC-II^pos^ only are able to suppress T cell proliferation under contact-dependent mechanisms. Regulatory neutrophils are highly enriched in lymphoid organs as compared to their MHC-II^neg^ counterparts and in the mouse they express PDL-1, an immune checkpoint involved in T-cell blockade. Our results emphasize neutrophils as true partners of the adaptive immune response, including in domestic species. They open the way for discovery of new biomarkers and therapeutic interventions to better control cattle diseases.

## INTRODUCTION

Neutrophils are major partners of the innate immune system and are considered the first line of defense against microbes[1]. They originate from the bone marrow (BM) and circulate in large numbers in the blood from where they are swiftly recruited to sites of inflammation or infection to fight danger. They are functionally equipped to rapidly phagocytose invading microbes via a variety of receptors[2] and rapidly deliver large amounts of Reactive Oxygen Species (ROS) into phagocytic vacuoles to kill microbes[3]. If not tightly controlled these dangerous weapons lead to tissue damage and neutrophils are often a signature of uncontrolled inflammation or infection.

Neutrophils are the dominant cell population circulating in blood in a wide range of animal species and humans[4]. However, in cattle or mouse blood they are less numerous than lymphocytes[5; 6]. Compared to human or mouse neutrophils, bovine neutrophils present peculiarities such as unique receptors or the lack of N-formylated chemotactic peptide receptor[5]. Neutrophil counts are an important signature of cattle condition. In lactating cows, they represent the most part of an abnormal somatic cell count in milk from mastitic cows. Indeed, their swift recruitment into the mammary gland is of critical importance in the fight against invasive pathogens[7]. For a long time, neutrophils were thought to only travel from their birth place, the BM, to blood and tissues in response to inflammatory signals or infections. However, this has recently been revisited by a number of authors who have identified neutrophils residing in spleen[8; 9], lymph nodes[10] or lung vasculature[11]. These new locations and features of resident neutrophils have been discovered in mice and humans but are not well documented in other animal species, including cattle, under normal conditions. Long considered as “suicide bombers”, neutrophils were recently upgraded as global players of the immune response along with the discovery of new functions[12]. They fully participate in shaping adaptive immunity by suppressing T cell proliferation and activity[13; 14] or by promoting IgM and IgG production by splenic B cells[9]. This broad range of phenotypes and functions was recently linked to discrete populations of neutrophils characterized by unbiased single cell analyses[15]. Like macrophages and dendritic cells, neutrophils are now recognized as plastic cells able to respond and adapt to their environment. Again, although neutrophil diversity and plasticity are now well documented for mouse and human neutrophils, they remain unknown for domestic species including cattle.

Here we conducted a parallel thorough characterization of neutrophils present in mouse BM or circulating in cattle blood at steady state. We define this term as animals without any apparent sign of disease or infection. In other words, animals where the immune system is at homeostasis. We discovered a new population of regulatory neutrophils that displayed suppressive activity on T-cells, at steady state.

## MATERIALS AND METHODS

### Animal protocols and sampling

Experimental protocols complied with French law (Décret: 2001–464 29/05/01) and European directive 2010/63/UE for the care and use of laboratory animals and were carried out under Authorization for Experimentation on Laboratory Animals Number D-37-175-3 (Animal facility UE-PFIE, INRA Centre Val de Loire for mice) and E 37-175-2 (UE-PAO, INRAE Centre Val de Loire for bovine). Animal protocols were approved by the “Val de Loire” Ethics Committee for Animal Experimentation and were registered to the French National Committee for Animal Experimentation under N°2016091610026164.V3 (mice) or N°2016040410531486 (cattle). Cattle remained in their environment (UE-PAO) and no animal was sacrificed for this work. Six-to eight-week-old C57BL/6 mice and OT-II transgenic mice for the OVA MHC class II complex-specific TCR[16] on a C57BL/6 genetic background were bred at the resident PFIE animal facility before use.

Bovine blood was collected from Holstein Friesian cows at the jugular vein into vacutainer K2 EDTA tubes. Broncho-tracheal lymph nodes, spleen biopsy and sternum BM were collected post-mortem at a commercial abattoir from Blonde d’Aquitaine, Limousine, Charolaise and Highlands cows. Mice were euthanized by CO2 inhalation. Blood (from heart), spleen, inguinal lymph nodes and femurs were collected.

### Preparation of cells

Tubes containing blood were centrifuged at 1000x*g* for 10 min at 20°C before removal of the plasma layer and buffy coat. In indicated experiments, buffy coat was used in proliferation assay (see below). Red Blood Lysis Buffer (Sigma 94 R7757) (4 vol/1 vol of blood) was added for 5 min at room temperature to lyse red blood cells. Cells were washed twice in D-PBS with 2mM EDTA. Mouse neutrophils were generally purified from the BM by positive magnetic selection with anti–Ly-6G PE-conjugated Ab (1A8; BD Biosciences) and anti-PE microbeads (Miltenyi Biotec) as described previously[17] except for cell sorting, where mouse neutrophils were enriched from BM by positive magnetic selection with anti–CD11b microbeads (Miltenyi Biotec) before labelling with antibodies. Lymph nodes (LN) and spleen biopsies were disrupted mechanically and cells were filtered through 100μm nylon cell strainer (BD Falcon), and washed twice. Cells were suspended in RPMI-1640 supplemented with 2mM L-Glutamine, 10mM HEPES and 1mg/mL of BSA with extremely low endotoxin level (≤1.0 EU/mg) (hereafter referred to as RPMI complete medium). Cell counts were determined after staining with Türk solution and numerated with a Malassez’s chamber.

### Flow cytometry

Bovine cells were suspended in PBS with 10% of horse serum (Gibco), 2mM EDTA and labeled for 30 min with primary antibodies (see Supplementary Table S1). Mouse cells were suspended in PBS 2mM EDTA. After saturation with anti-CD32/CD16, cells were incubated 30min with fluorescent mAb. After washes in D-PBS (300x*g*, 10 min, 4°C), cells were labeled 30min with the corresponding fluorescent-conjugated secondary antibodies. Bovine cells were washed and fixed with BD cell Fix diluted 4 times in PBS. Data were acquired with a LSR Fortessa™ X-20 Flow cytometer (Becton Dickinson) and results analyzed with Kaluza software (Beckman Coulter).

For neutrophil subsets purification, cell concentrations were adjusted to 10^7^cells/mL and sorted with a MoFlo Astrios^EQ^ high speed cell sorter (Beckman Coulter) according to our previously published protocol [18]. Sorted cells were spread on microscope slides (Superfrost, Thermo) by cytocentrifugation (3 min, 700x*g*) and stained with May-Grünwald and Giemsa with the RAL 555 kit (RAL diagnostics).

### Neutrophils functional tests

Sorted neutrophils viability was evaluated after 1 h stimulation with 100ng/mL of LPS (*E.coli* 0111:B4, Sigma). Neutrophils were then incubated 15min at room temperature with anti-annexinV antibodies in binding buffer (BD Bioscience). Cells were washed and incubated 15min at room temperature with streptavidin-APC-Cy7 (BD Bioscience). Cells were washed and incubated 5 min with 20µg/mL of propidium iodide (BD Bioscience) and then directly analyzed with LSR Fortessa X-20 flow cytometer. Phagocytosis was measured using pHrodo™ Red *E. coli* BioParticles® Conjugated (MolecularProbes®) following the manufacturer’s instructions. Briefly, purified neutrophils were first incubated in a 96 wells microplate for 30 min at 37°C in RPMI complete medium with or without 2µg/well of cytochalasin D (Sigma), and then for 1 h with 20µg/wells of pHrodo *E. coli* BioParticles. Fluorescence was directly measured with the LSR Fortessa™ X-20 Flow cytometer. ROS produced by neutrophils were quantified using the CellROX® Orange Flow Cytometry Assay Kits (MolecularProbes®, C10493) following the manufacturer’s instructions. Briefly, purified neutrophils were first incubated for 1 h at 37°C in RPMI complete medium with or without 400µM of TBHP in a 96-wells black microplate and then for 30 min with 100nM CellROX®. Fluorescence was measured with the LSR Fortessa™ X-20 Flow cytometer. To test the bacterial killing capacity of neutrophils, we used the *Escherichia coli* P4 strain, isolated from a clinical case of bovine mastitis which causes severe infections in mice[19]. *E. coli* P4 bacteria were grown in 10 mL BHI medium overnight at 37°C without agitation. Bacteria were then diluted in BHI medium (1 vol/100 vol) and incubated for 6 h at 37°C without agitation. Bacterial concentration was determined by the optical density at 600 nm and adjusted at 4.10^5^ CFU/mL in RPMI complete medium. Purified neutrophils were infected at a MOI of 0.2 in RPMI complete medium in 1.5mL Eppendorf tubes for 90 min at 37°C under agitation on a Rotator SB3 (Stuart equipment). Bacteria in RPMI complete medium without neutrophils were used as the reference to asses killing. Bacterial dilutions (in PBS 0.25% SDS) were plated on TSA agar plates and incubated for 16 h at 37°C before numeration of CFUs.

### RNA extraction and gene expression analysis

Total RNAs were extracted from cell-sorter purified cells using NucleoSpin RNA kit with a DNase treatment (Macherey Nagel) and reverse transcribed with iScript™ Reverse Transcriptase mix (Biorad) according to the manufacturer’s instructions. Primers (Eurogenetec) are listed in Supplementary Table S2. Primers validation was performed on a serial diluted pool of cDNA (a mix of cDNA from spleen, lung, LN, blood and BM cells for both species) with a LightCycler® 480 Real-Time PCR System (Roche). Gene expression was then assessed with the BioMark HD (Fluidigm) in a 48×48 wells plate, according to the manufacturer’s instructions. The annealing temperature was 60 and 62°C for bovine and mouse samples respectively. Data were analyzed with Fluidigm RealTime PCR software to determine the cycle threshold (Ct) values. Messenger RNA (mRNA) expression was normalized to the mean expression of three housekeeping genes for each animal species to obtain the ΔCt value. Principal Component Analysis (PCA) and hierarchical clustering were performed with ΔCt values in R studio (Version 1.3.959, © 2009-2020 RStudio, PBC), using respectively the FactoMineR and pheatmap packages. Ward’s minimum variance method was applied for clustering, with dissimilarities squared before clustering (ward.D2). ΔCt values were centered to the median for clustering.

### Measure of T-cell suppressive activity of neutrophils

For the mouse, splenocytes from OT-II mice were collected, homogenized to single-cell suspensions through nylon screens and resuspended in RPMI medium (Gibco) supplemented with 10% decomplemented fetal bovine serum (Gibco), 2 mM L-glutamine (Gibco), 100 U penicillin and 100 μg/ml streptomycin (Gibco). 10^5^ cells/well were distributed in a 96-wells round bottom plate (BD Falcon). OT-II splenocytes proliferation was induced by addition of 2 μg/ml of the OVA peptide 323-339 (Polypeptide Group). As indicated, purified neutrophils were added to the culture at a ratio of 1 neutrophil:10 splenocytes in a final volume of 200µL. Wells without neutrophils were used as reference for maximal proliferation.

As indicated, neutrophils were separated from splenocytes by placing them in a HTS Transwell-96 permeable device with 0.4 μm pore and polycarbonate membrane and adapted receiver plate (Corning, reference CLS3381). To test the role of MHC-II and CD11b molecules in the suppression mechanism, neutrophils were also treated 1 h before incubation with splenocytes with 15µg/mL anti-CD11b mAb (clone M1/70) or rat IgG2bκ as isotype control ; anti MHC-II mAb (clone 2G9) or rat IgG2aκ as isotype control. Plates were incubated at 37°C with 5%CO_2_. Cell proliferation was quantified after 3 days of culture using CyQUANT Cell Proliferation Assay tests (ThermoFisher) according to the manufacturers’ instructions.

For the bovine, a Mixed Lymphocyte Reaction assay was set up by mixing blood cells from two genetically distant cows with cow N°1 as the responding animal and cow N°2 as the stimulating animal. Briefly, blood was centrifuged at 1000x*g*, 15min, 20°C and buffy coats were collected and diluted 4 times in PBS. PBMCs were collected at the interface of 1.077 density Percoll gradient (GE Healthcare) after centrifugation at 400*g*, 15min, 20°C, without brake. PBMCs from cow n°1 (responding) remained untreated while PBMCs from cow n°2 (stimulating) were incubated for 30 min at 38.5°C in 5% CO2 with 50µg/mL of mitomycin C from *Streptomyces caespitosus* (Sigma, M4287), to block their proliferation. After three washes, 10^5^ stimulating cells were mixed with 10^5^ responding cells in 96 wells plates (ratio 1:1) in a total volume of 150µL in RPMI complete medium supplemented with 100U/mL penicillin and 100μg/ml streptomycin. The negative control was responding cells alone and the reference maximal proliferation was responding and simulating cells together at ratio 1:1. Proliferation that started at day 6 was stopped at day 9. As indicated, at day 4, 10^5^ of purified syngenic neutrophils from the responding animal were added to the well in a final volume of 200 µL. In one experiment, neutrophils were placed in a HTS Transwell-96 permeable device as for the mouse system. To quantify proliferation, plates were centrifuged at day 6 and day 9 for 5 min at 300x*g*, supernatants were discarded and cells were frozen at -80°C before addition of reagents from the CyQUANT Cell Proliferation Assay kit (ThermoFisher) as indicated by the manufacturer.

### Statistical analysis

Individual data and the median were presented in the Figures. Statistical analyses were performed with Prism 6.0 software (GraphPad). Analyzes were performed on data from 2 to 6 independent experiments, Mann Whitney non-parametric tests or 2way ANOVA test were used. Represented p-values were: \**p* < 0.05; \*\**p* < 0.01, and \*\*\**p* < 0.001.

## SUPPLEMENTARY INFORMATION

Supplemental information can be found with this article online. Should the reader need additional details, please email a request to corresponding authors.

## RESULTS

### Neutrophils represent discrete populations in mouse and cattle

In cattle, neutrophils are often isolated by simple centrifugation of freshly collected blood to separate them from the buffy coat. They segregate at the bottom of the tube with red blood cells[20; 21]. In mice, because of very small volumes, blood sampling is less practical. Therefore, high numbers of neutrophils are directly extracted from the BM by magnetic selection following labelling with antibody against Ly-6G which is highly expressed by mouse neutrophils[22]. These methods, that lead to a fair level of purity for neutrophils, are convenient for most assays because they are quick and preserve these fragile cells. We prepared cattle and mouse neutrophils using these rapid procedures and analyzed them by flow cytometry (Fig. 1). Following mouse bone-marrow cell preparation and labeling with anti-Ly-6G and magnetic separation (Fig. 1A), banded-cells of apparent homogeneity were observed after May-Grümwald-Giemsa (MGG) staining (Fig. 1B). However, the SSC and FSC profile by flow cytometry displayed two discrete populations of heterogeneous size. Moreover, double labelling of these “pure” neutrophils with anti-Ly-6G and anti-CD11b distinguished CD11b^hi^ from CD11b^med^ cell populations (Fig. 1C). In cattle, after elimination of the buffy coat containing PBMCs, cells were collected from lower 2/3 of the tube (Fig. 1D) and analyzed after centrifugation and MGG staining. This revealed heterogeneity of this cell fraction with the presence of eosinophils in variable proportions (Fig. 1E). After labelling with anti G1, a marker that is highly expressed on the surface of bovine neutrophils[23; 24] and anti CD11b, cells were analyzed by flow cytometry. After SSC and FSC gating on the granulocytes populations (Fig. 1F), we distinguished CD11b^pos^ G1^low^ eosinophils (representing between 2 and 8% of the SSC x FSC granulocyte gate depending on the animal) from G1^hi^ neutrophils. Among G1^hi^ neutrophils CD11b labelling segregated two populations: a main population of CD11b^med^ neutrophils (around 87% of the granulocyte gate) and a minor population (around 1,5%) of CD11b^hi^ neutrophils (Fig. 1F).

**Figure 1.**
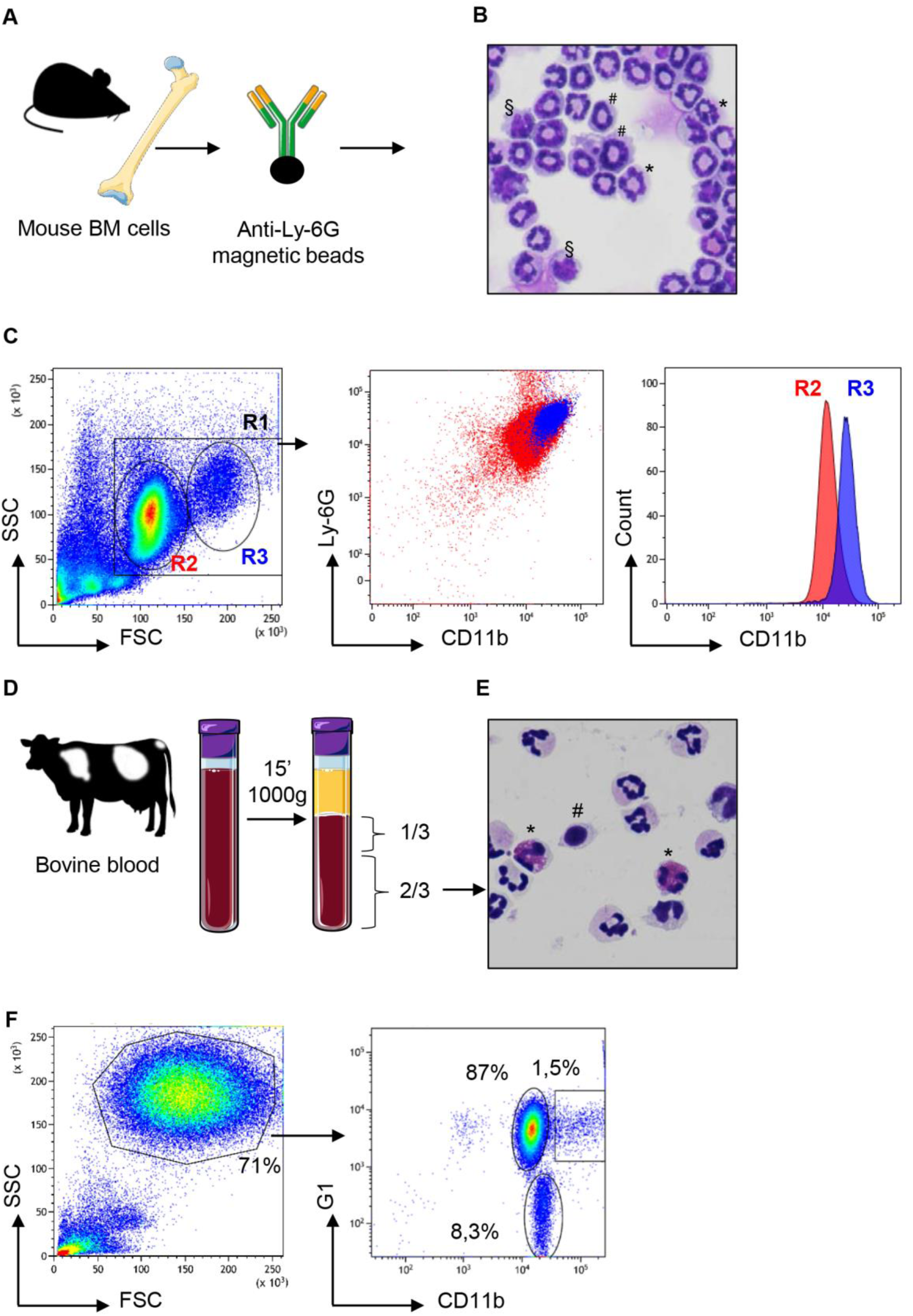
Neutrophils from mouse bone marrow or cattle blood display heterogeneous profiles. (**A**) BM cells from C57BL/6 adult mice were magnetically purified with anti-Ly6G beads (**B**) These cells stained with May Grünwald Giemsa after cytocentrifugation on glass slides displayed heterogenous profiles with segmented nuclei (*), banded nuclei (#) as well as some nuclei with a mononuclear kidney shape (§). (**C**) After anti-Ly-6G magnetic purification, mouse cells were analyzed by flow cytometry using the LSR Fortessa™ X-20 apparatus and in the “granulocytes” R1 gate, a bimodal distribution of FSC^med^ (R2) and FSC^high^ (R3) cells was observed on a linear scale (first panel). Labelling with anti-Ly6-G and anti-CD11b confirmed this heterogeneity after dot plot analysis (middle panel) with bimodal distribution of CD11b^med^ (R2) and CD11b^hi^ (R3) cells as displayed on the histogram analysis (right panel). (**D**) Blood from Hostein Friesian cows was centrifuged and the lower 2/3 of the tube enriched in neutrophils were collected. (**E**) These cells, as observed after cytocentrifugation and staining by May Grünwald Giemsa, were mainly neutrophils, characterized by their polylobed nucleus but monocytes (#) and eosinophils (*) were also observed in various proportions depending on the animal. (**F**) These cells were analyzed by flow cytometry and FSC x SSC dot plot showed granulocytes in a gate that represented 71% of analyzed cells (left panel). After labeling with anti-G1 and anti CD11b, dot-plot analysis of the “granulocytes” gate (right panel) revealed three subsets: eosinophils that were negative for the G1 marker (around 8% of granulocytes) and two subsets of G1^pos^ neutrophils that were CD11b^med^ (87%) or CD11b^hi^ (1.5%). The most representative animal is shown (n=4 for mouse, n=6 for bovine), although in cattle the proportion of eosinophils and CD11b^hi^ neutrophils circulating in blood varied between animals.

As we were intrigued by the presence of different neutrophil populations in mouse BM and cattle blood at steady state, we performed a more thorough characterization by flow cytometry using a panel of markers (Fig. 2). In mouse, CD11b was used to gate myeloid cells (Fig. 2A) and a combination of anti-Ly-6C and anti-Ly-6G allowed us to distinguish *bona-fide* neutrophils (Ly-6G^hi^) from monocytes (Ly-6G^neg^)[25]. Among Ly-6C^+^ Ly-6G^hi^ cells, a minor subset (1,2%) of MHC-II^pos^ Ly-6C^+^ Ly-6G^hi^ neutrophils was clearly distinguished from the main population (92,5%) of the MHC-II^neg^ neutrophils. In cattle, among the neutrophils that highly expressed the G1 marker, MHC-II^neg^ neutrophils represented the main population (93%). However, a minor population of G1^hi^ neutrophils (around 1,5%) expressed MHC-II antigens on the surface as observed in the mouse (Fig. 2A). The two MHC^neg^ et MHC^pos^ subsets were sorted by flow cytometry in both species. After centrifugation onto glass slides and MGG staining, they were indistinguishable under a microscope (Fig. 2B). Sorted MHC-II^neg^ and MHC-II^pos^ neutrophils were then labelled with a panel of antibodies (Fig. 2C). In both bovine and mouse, in comparison with classical MCH-II^neg^ neutrophils, the MHC-II^pos^ subset overexpressed CD11b on the surface as well as L-selectin CD62L. The CD14 LPS coreceptor was not detected on MHC-II^neg^ neutrophils. By contrast around 50% of MHC-II^pos^ neutrophils displayed this receptor on their surface in both the mouse and the bovine. In mouse, we also observed that the MHC-II^pos^ subset expressed high levels of CD44 as well as CD274 (PDL-1) the ligand for PD-1 involved in T cell exhaustion[26]. Since neutrophils are fragile cells which could be exacerbated by flow cytometry sorting, we next asked if the two subsets displayed different *ex-vivo* survival times. The two subsets were incubated with LPS for 1 hour and stained with annexin V and propidium iodide (Fig. 2D). The large majority of neutrophils remained alive after this treatment and no significant difference was observed between classical MHC-II^neg^ neutrophils (97% alive in mouse or bovine) and MHC-II^pos^ neutrophils (95% alive in mouse or bovine).

**Figure 2.**
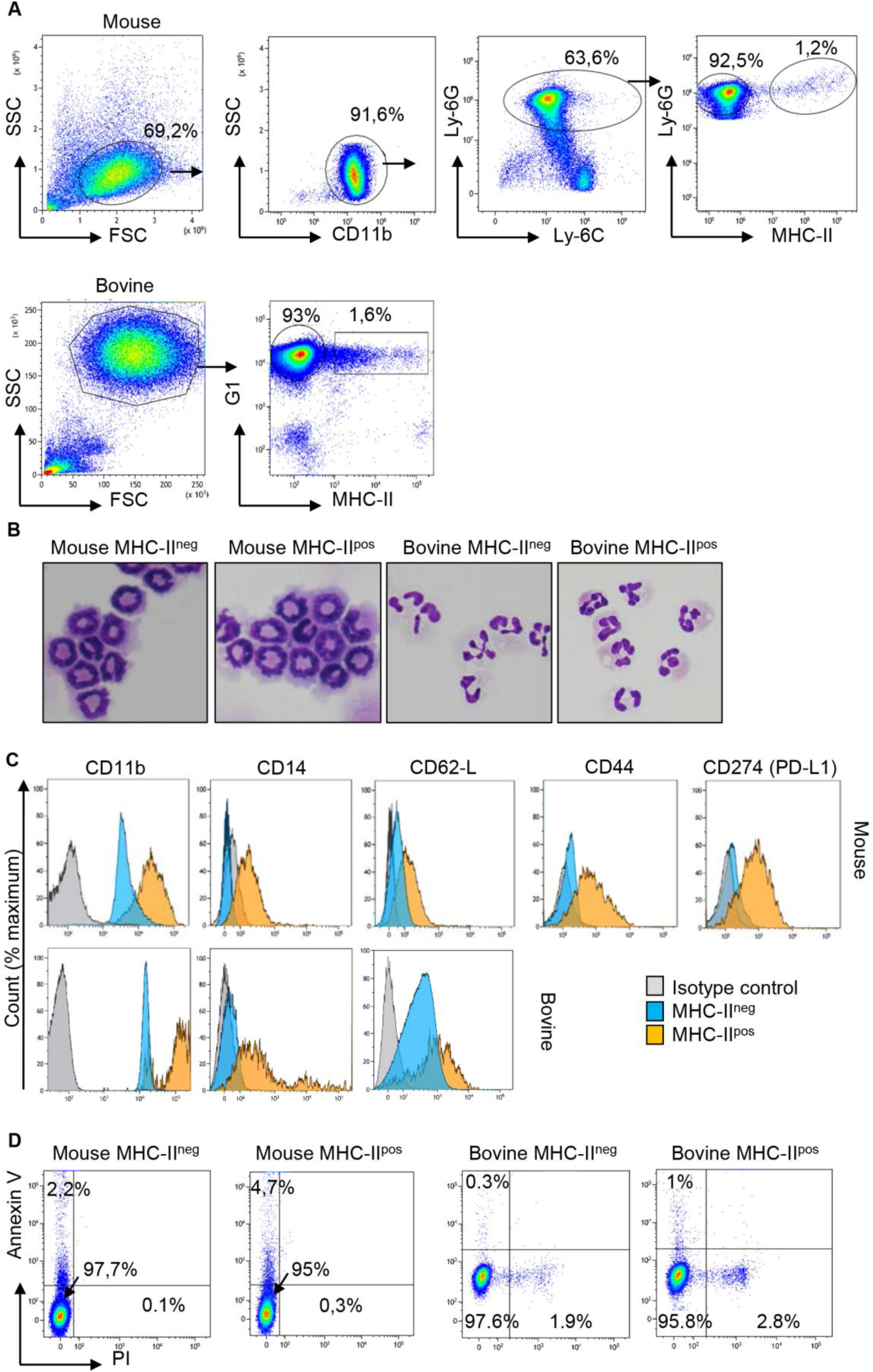
Two populations of neutrophils can be distinguished on the basis of specific surface makers, including MHC-II, in the mouse and the bovine. **(A)** Cells were prepared from mouse BM or cattle blood as described in Fig 1 and neutrophils were sorted with a MoFlo Astrios^EQ^ apparatus after labelling with eFluor viability dye 780, anti-CD11b, anti-Ly-6C, anti-Ly-6G and anti MHC-II in the mouse or eFluor viability dye 780, anti-G1 and anti-MHC-II in the bovine. Representative plots that show the major proportion of MHC-II^neg^ and the minor proportion of MHC-II^pos^ neutrophils in both species are depicted. **(B)** The two MHC-II^neg^ and MHC-II^pos^ neutrophil populations among total CD11b^pos^ Ly6-G^pos^ Ly6-C^pos^ mouse neutrophils, or G1^pos^ bovine neutrophils, were sorted by flow cytometry (purity >99%), cytocentrifuged and were indistinguishable after May Grünwald Giemsa staining. **(C)** Histograms represent expression of CD11b, CD14, CD62-L for both species and CD44 and CD274 /PD-L1 for the mouse samples on the surface of sorted MHC-II^neg^ and MHC-II^pos^ neutrophils in comparison to isotype controls. **(A-C)** Results from one representative animal are depicted (n=6 for mouse, n=6 for bovine). (**D**) After purification by flow cytometry, MHC-II^neg^ and MHC-II^pos^ neutrophils from the two species were incubated with LPS for 1 h and labelled with annexin-V and propidium iodide (PI) to analyze apoptotic (annexin V+ ; PI-) and dead cells (annexin V+ ; PI+). Dot plots from one representative animal are depicted (n=4 for mouse, n=3 for bovine).

Therefore, in mouse BM or in cattle blood, two subsets of neutrophils could be distinguished at steady state by MHC-II labelling. Both subsets displayed similar apoptosis and death profiles *ex-vivo* after LPS stimulation.

### MHC-II^pos^ neutrophils are enriched in lymphoid organs

Although neutrophils are mostly present in the BM reservoir and in circulating blood, recent data in the mouse model and in humans highlighted their wider distribution in tissues[27] including lymphoid organs. We next examined the distribution of MHC-II^neg^ versus MHC-II^pos^ neutrophils in different compartments in both mouse and cattle. Cells were extracted and prepared from BM, lymph nodes (LN, inguinal in the mouse and tracheobronchial in bovine) and spleen. For analysis of circulating neutrophils, total blood leukocytes were analyzed by flow cytometry (see Supplemental Fig. S1 for gating strategy) after lysis of red cells. In the BM, neutrophils represented around 30% of total leukocytes in both species (Fig. 3A). This was expected since BM is the principal compartment hosting neutrophils. In blood, neutrophils represented 30% or 20 % of total cells in mouse and cattle, respectively. In both the mouse and the bovine LN, even though neutrophils represented less than 1% of cells at steady state, they were consistently detected. Similarly, between 2 and 3% of cells collected from mouse or bovine spleen after lysis of red blood cells were neutrophils (Fig. 3A). In all analyzed compartments classical neutrophils as well as MHC-II^pos^ neutrophils were detected (Fig. 3B). Interestingly, whereas MHC-II^pos^ neutrophils represented only 2,4 ± 0,6 % or 4,6 ± 1,4 % of total neutrophils in the mouse bone-marrow or cattle blood, respectively, they represented around 40% or 60% of total neutrophils in mouse or bovine lymph nodes, respectively. In spleen, around 20% of total mouse neutrophils and 30% of bovine were MHC-II^pos^ (Fig. 3B). Therefore MHC-II^pos^ neutrophils were highly enriched in lymphoid organs in both species, at steady state.

**Figure 3.**
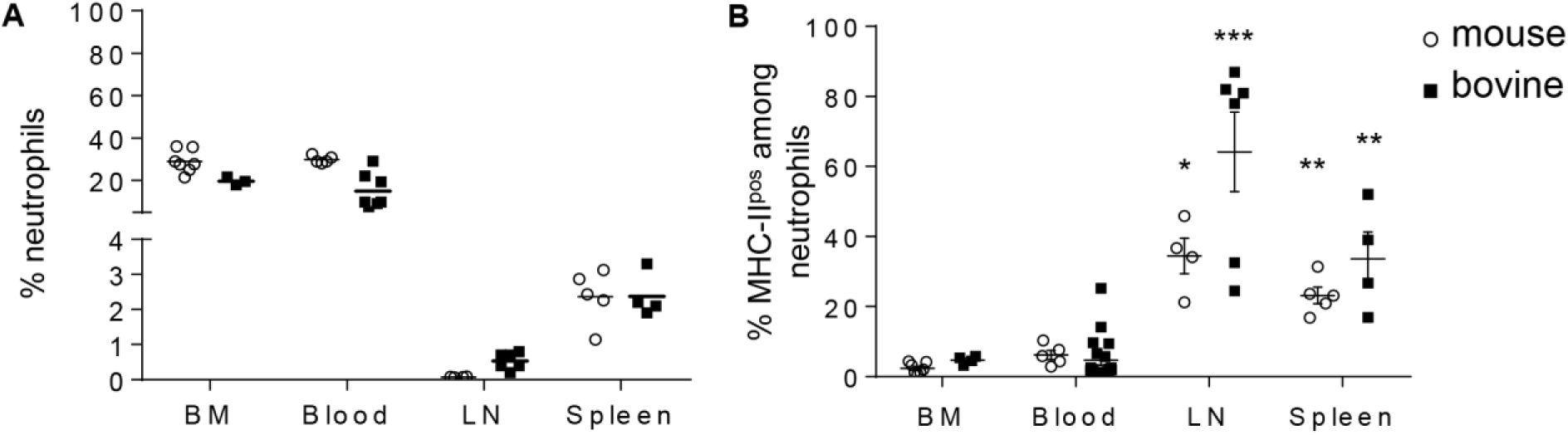
MHC-II^pos^ neutrophils are enriched in lymphoid organs in both species. **(A-B)** Cells were prepared from BM or blood from the two species and neutrophils analyzed by flow cytometry as in Fig 2. Neutrophils present in the lymphoid organs were also analyzed after collecting post mortem the inguinal (mouse) or tracheobronchial (cattle) lymph nodes (LN) and spleen. (**A**) The percentage of Ly6-G^pos^, Ly6-C^pos^ mouse neutrophils or G1^pos^ cattle neutrophils among total leukocytes and (**B**) The percentage of the MHC-II^pos^ subset among total neutrophils, was analyzed in each compartment. Individual data and the median in each group are presented. This percentage was significantly higher in lymphoid organs than in mouse BM or cattle blood. * *P*<0.05; **, *P*<0.01; ***, *P*<0.001 (Mann Whitney non-parametric test).

### RNA profiling distinguishes classical from MHC-II^pos^ neutrophils in both the mouse and the bovine

To analyze the transcriptional profile of MHC-II^pos^ and MHC-II^neg^ neutrophils, we sorted the two populations from either BM (mouse) or blood (cattle) to more than 99% purity and performed a transcriptomic profiling using 48 validated primer pairs (Table S1) designed to cover a large set of neutrophils functions such as synthesis of cytokines or chemokines, enzymes stored in granules, surface receptors, as well as transcription factors. Some weakly expressed genes were removed from the panel (highlighted in Table S1). We then performed unsupervised Principal Component Analysis (PCA) and observed that the different subsets were significantly discriminated (Fig. 4A). The first axis of the PCA (explaining 26.4% of the variance) separated bovine and mouse samples, as expected. The second axis (explaining 19.1% of the variance) clearly separated the MHC-II^pos^ from the MHC-II^neg^ neutrophils in each species (Fig. 4A), indicating that they belonged to different subsets. As expected, in the bovine, gene expression profiles were more dispersed than in the inbred mouse, due to inter-individual variability. In both species, the two neutrophil subsets were also segregated by hierarchical clustering based on ΔCT values (Fig. 4B). Most of the genes were significantly less expressed in MHC-II^pos^ neutrophils as compared to MHC-II^neg^. However, expression of the following genes was significantly higher in the MHC-II^pos^ subset: *elane, proteinase3, PD-L1* in murine and *MHC-II*; *AHR* in bovine cells. A trend for higher expression was observed in MHC-II^pos^ neutrophils for *CD14* (both species) and *proteinase3* (bovine). Therefore, in both species the transcriptomic signature significantly differed between the two neutrophil subsets which suggested different biological roles.

**Figure 4.**
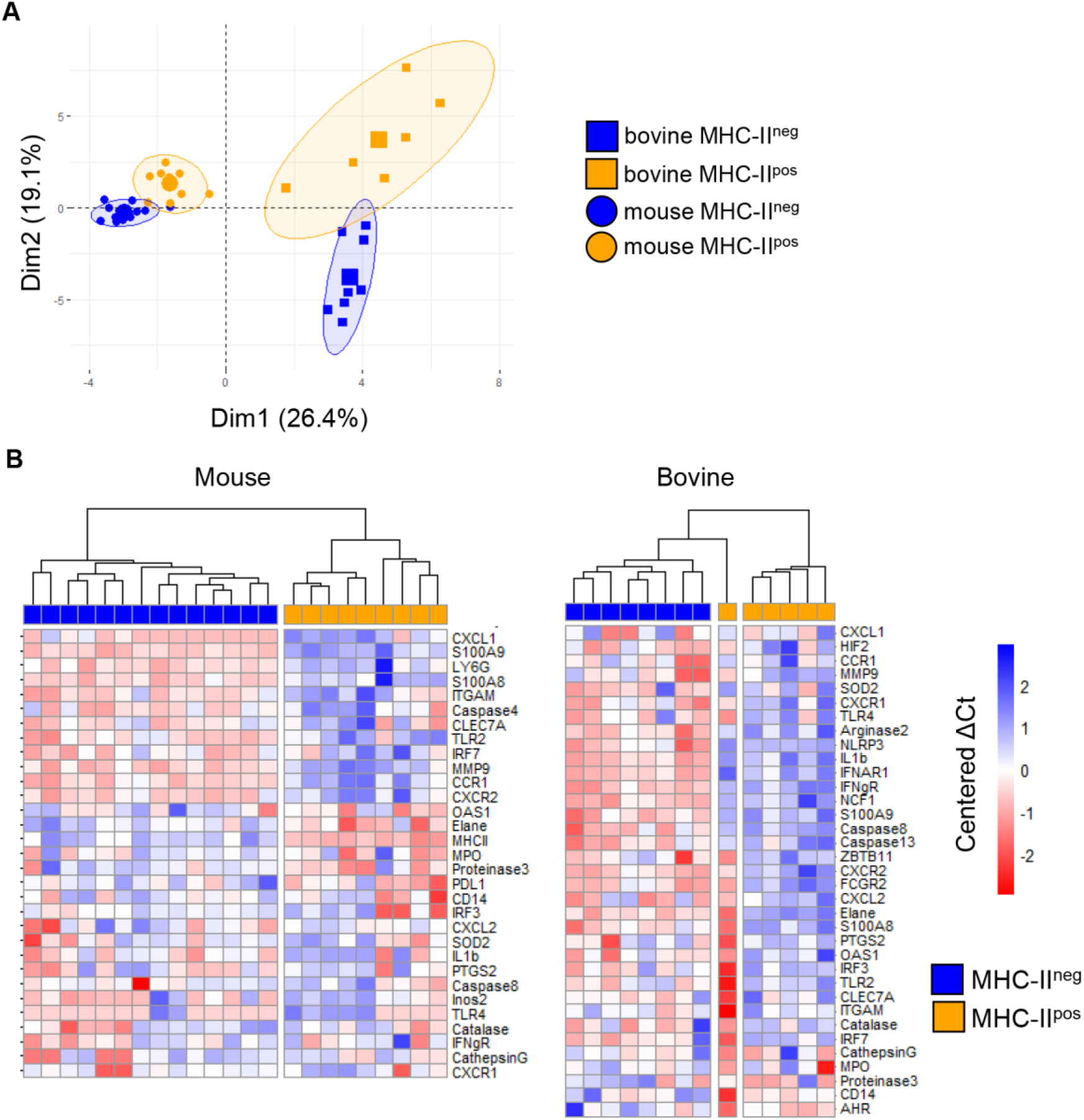
MHC-II^pos^ and MHC-II^neg^ neutrophils display distinct transcriptomic signatures. Gene expression of the two purified neutrophil subsets (more than 99% purity) from the two species was assessed by quantitative Real Time PCR using sets of primer pairs designed to cover a large range of neutrophils functions. mRNA expression was normalized to the expression of three housekeeping genes for both species to calculate the ΔCt values (**A)**. Principal Component Analysis (PCA) was performed on ΔCt values and the two first dimensions of the PCA plot are depicted. Inclusion in groups of sets of samples are delineated in the colored areas on the graph with a confidence level of 90% (**B)** Hierarchical clustering of gene expression was performed on median centered ΔCt values for mouse and bovine data sets, using the ward.D2 method. For each gene, ΔCt values were centered to the median ΔCt value. Higher or lower ΔCt expression compaired to the median value were represented respectively in deep to light red or blue. Expression of a selected set of genes indicated on the figure was clearly distinct in MHC-II^pos^ neutrophils or MHC-II^neg^ neutrophils. Data represent individual samples (Mouse: n= 14 for MHC-II^neg^, n=9 for MHC-II^pos^ Bovine: n=8 for MHC-II^neg^ n=6 for bovine MHC-II^pos^).

### MHC-II^pos^ neutrophils produce higher levels of ROS than MHC-II^neg^ but similarly phagocytose bioparticles and kill bacteria

In order to gain further insight into the functions of the two neutrophil populations, we first compared their phagocytic ability using conjugated pHrodo(tm) Red *E. coli* BioParticles*(tm)* that only fluoresce once inside the phagosome or endosome of the cell[28]. Neutrophils were then analyzed by flow cytometry and the percentage of phagocytosis was directly correlated with the mean fluorescence intensity (see supplementary Fig. S2). Phagocytosis by MHC-II^neg^ or MHC-II^pos^ neutrophils was compared in individual animals from each species. Both subsets actively phagocytosed bioparticles which was dramatically reduced by treatment with cytochalasin D (Fig. 5A). We did not detect a significant difference in phagocytosis between MHC-II^neg^ and MHC-II^pos^ neutrophils neither in the mouse nor the bovine (Fig. 5A). We next measured their potential for total ROS production after incubation with the non-specific chemical inducer *tert-butyl hydroperoxide* (TBHP) using the fluorescent probe CellROX that fluoresces when oxidized (see supplementary Fig. S2 for gating strategy). Mouse bone-marrow neutrophils incubated with medium produced very low levels of ROS whereas bovine blood neutrophils produced higher levels (Fig. 5B). As expected, TBHP treatment dramatically increased ROS production by both MHC-II^neg^ and MHC-II^pos^ neutrophils in both species. However, in both conditions and for both species, MHC-II^pos^ neutrophils produced significantly more ROS than MHC-II^neg^ cells (Fig. 5B). We next analyzed the killing activity, under non opsonic conditions, of the two subsets against the *Escherichia coli* P4 strain isolated from a case of bovine clinical mastitis[19]. Both types of neutrophils efficiently killed *E. coli* and no significant difference was observed between MHC-II^neg^ or MHC-II^pos^ neutrophils (Fig. 5C).

**Figure 5.**
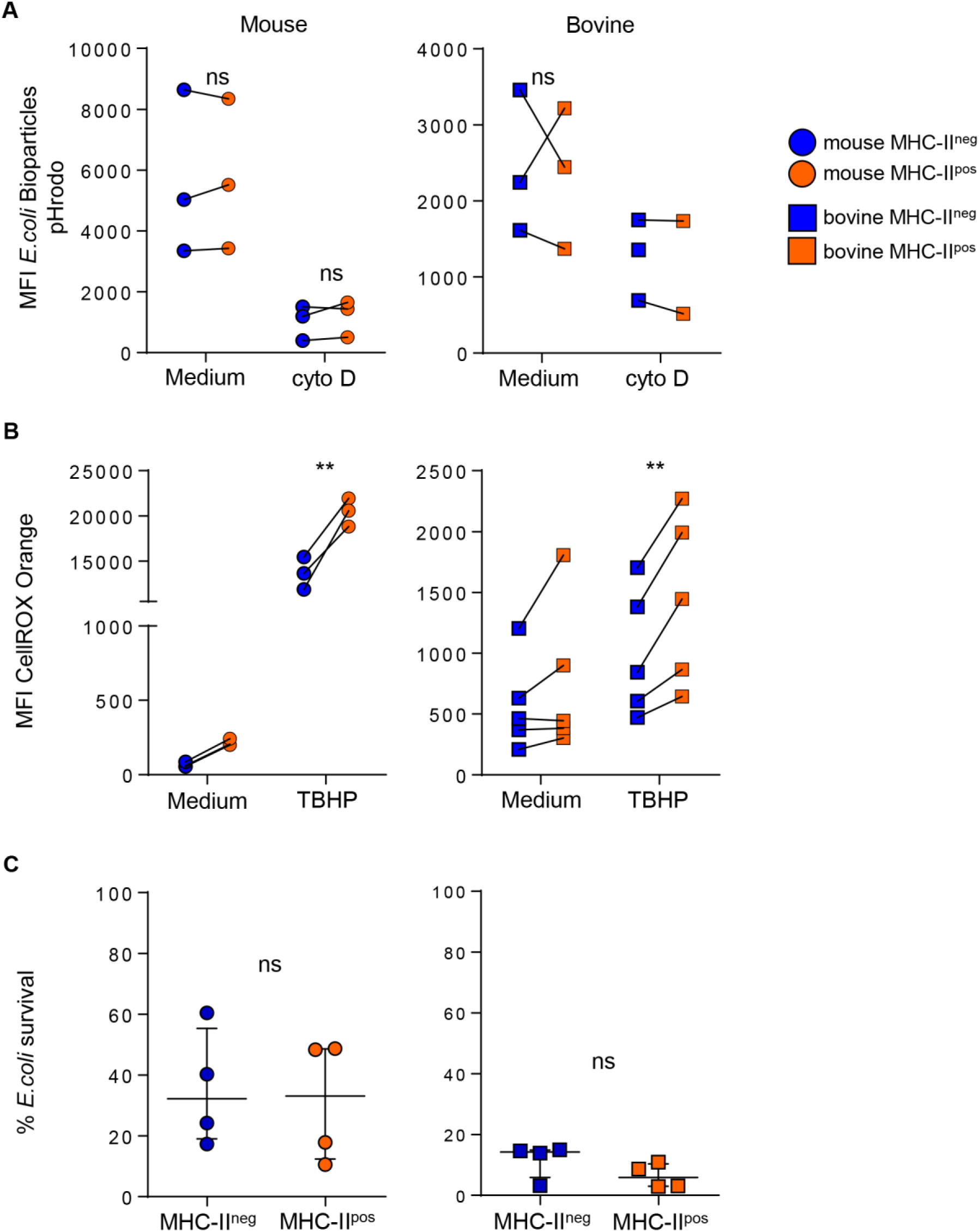
MHC-II^pos^ phagocytose bioparticles and kill *E. coli* similarly to MHC-II^neg^ neutrophils but produce higher levels of ROS. (**A)**. After purification by cell sorting from the BM (mouse) or blood (bovine), phagocytosis by MHC-II^pos^ or MHC-II^neg^ neutrophils was assessed using pHrodo *E.coli* bioparticles. Mean fluorescence intensity was directly correlated to ingested particles. Cells were treated with cytochalasin D as indicated. (**B)**. Oxidative stress was measured in MHC-II^pos^ and MHC-II^neg^ neutrophils using the CellROX Orange probe that reacts with all ROS species. Cells were activated with TBHP or incubated with medium alone and levels of ROS were measured by flow cytometry. **(A-B)** Mean fluorescence intensity in each sample is depicted and paired MHC-II^pos^ and MHC-II^neg^ samples were analyzed for each animal. (**C)**. Purified MHC-II^pos^ and MHC-II^neg^ neutrophils were infected with the *E. coli* P4 strain and bacterial survival was calculated by determining the ratio of bacteria incubated alone or in presence of neutrophils. Data represent n=4 independent experiments with neutrophils pooled from independent lots of 10 mice, or prepared from blood of n=4 independent cows. **(A-C)** *, *P*<0.05; **, *P*<0.01; ***, *P*<0.001 (2way ANOVA).

### MHC-II^pos^ but not MHC-II^neg^ neutrophils exert contact-dependent suppression of T cells at steady state

We wondered if surface proteins such as MHC-II (mouse and cattle) or PDL-1 (mouse) or enrichment of MHC-II^pos^ neutrophils in lymphoid organs could be linked to regulatory functions on T cells. To address this, we set up *in vitro* assays that were either antigen specific in the mouse (Fig. 6A) or polyclonal in cattle (Fig. 6C). Using the OT-II transgenic mice that bear the OVA peptide 323-339 - MHC class II complex-specific TCR[16] we observed strong proliferation of splenocytes when stimulated with OVA peptide for 72 hours. Taking this condition as the maximum proliferation (100%) we compared the impact of adding MHC-II^neg^ and MHC-II^pos^ neutrophils to the proliferating cells (ratio 10 splenocytes:1 neutrophil). The classical MHC-II^neg^ neutrophils had no measurable effect on OT-II splenocytes proliferation (Fig. 6B). By contrast, addition of MHC-II^pos^ neutrophils purified either from mouse BM or blood, decreased the capacity of CD4 OT-II cells to proliferate by 66 ± 3 % (Fig. 6B). The suppressive activity of mouse MHC-II^pos^ neutrophils depended on contact with the proliferating T-cells as no effect was observed when cells were separated by a Transwell device (Fig. 6B). We then used blocking antibodies in order to define if MHC-II or CD11b, that are both highly expressed on the surface of MHC-II^pos^ neutrophils, were involved in this suppressive activity. Both anti-MHC-II and anti-CD11b partially relieved suppression by MHC-II^pos^ neutrophils that reached 70 ± 7 and 75 ± 1% of total proliferation respectively (Fig. 6B). This indicated that both molecules were involved in the suppression mechanism although they were probably not the only ones.

**Figure 6.**
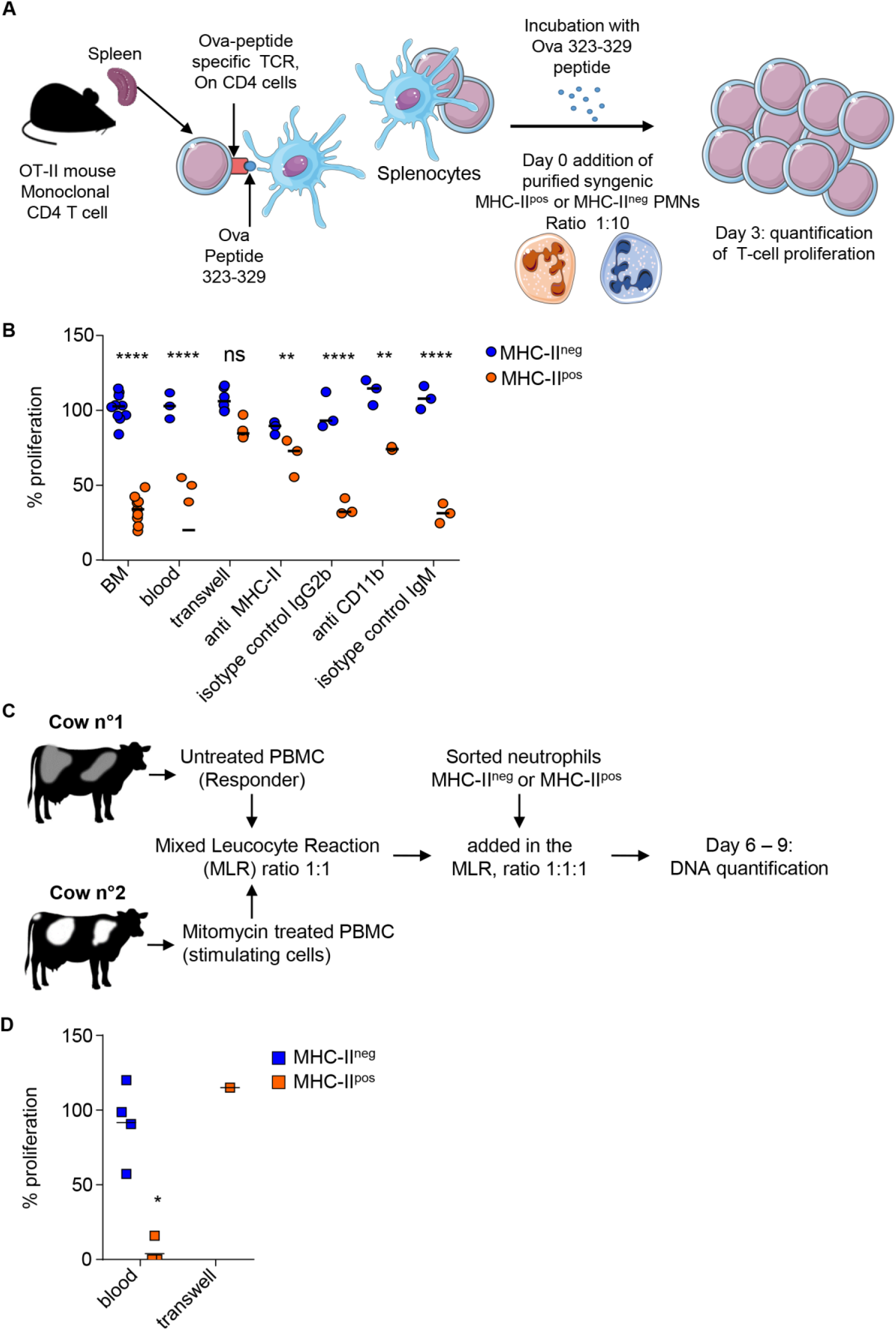
MHC-II^pos^ but not MHC-II^neg^ neutrophils exert suppression of T cells at steady state. **(A)** In the mouse an antigen -specific proliferation assay was set up with OT-II transgenic mice that carry the monoclonal population of T-cells specific for the Ova 323-329 peptide. Proliferation of splenocytes from OT-II mice activated with the Ova 323-329 peptide was set up as the maximum and compared to conditions where MHC-II^pos^ or MHC-II^neg^ neutrophils purified from syngenic C57BL/6 mice after cell sorting (99% purity) were added with a ratio of 1:10. (**B)** The percentage of T cell proliferation after addition of MHC-II^pos^ or MHC-II^neg^ neutrophils was calculated based on OT-II splenocytes proliferation with Ova peptide only. Neutrophils were prepared from the BM of syngenic mice except for the “blood” sample as indicated on the graph. Different conditions for the assay are also indicated, with cells separated by a transwell device, or neutrophils incubated with anti-MHC-II, anti-CD11b or isotype controls before addition to the proliferating splenocytes. **(C)**. For the bovine, a Mixed Leucocyte Reaction (MLR) was set up to assess polyclonal T cell proliferation. PBMCs from the responder animal were isolated and left untreated, while PBMCs from the stimulating animal were incubated with mitomycin C. PBMCs from the two cows were incubated at ratio of 1:1. To assess the impact of neutrophils on cattle T cell proliferation, 10^5^ of sorted MHC-II^pos^ or MHC-II^neg^ neutrophils from the responder animal were added to the reaction at day 4. DNA was quantified at day 6 and 9. **(D)** The proliferation was calculated by subtracting DNA values at day 9 from day 6 values. Proliferation measured in the MLR without neutrophils was defined as the reference 100% proliferation. The proliferation observed in the presence of sorted MHC-II^pos^ or MHC-II^neg^ neutrophils was calculated according to the DNA content (Day 9 – Day 6) and expressed as percentage of the reference value. In one experiment, MHC-II^pos^ neutrophils were separated from proliferating cells by a Transwell device. (**B and D)** Individual data and the median in each group are represented (3 independent experiments, n≥3 pool of 10 mice, n=4 cows, n=1 cow for transwell experiment; each represented value is the mean of technical triplicates). *, *P*<0.05; **, *P*<0.01; ***, *P*<0.001 Mann Whitney non parametric test).

In cattle, we set up a Mixed-Leukocyte-Reaction (MLR) by mixing PBMCs from a responder animal with mitomycin-C treated PBMCs from a genetically unmatched animal as stimulating cells (ratio 1:1) for a total of 9 days. After initial decline of total DNA content during 6 days due to cells dying in the wells, proliferation of T-cells from the responder animal was measured by an increase of the DNA content at day 9 in the control wells, correlating with polyclonal activation of T-cells (Fig. S3). Level of proliferation obtained under these control conditions was set as 100% (Fig. 6D). Addition of classical MHC-II^neg^ bovine neutrophils to the PBMCs (ratio 1:1) from the responder animal did not change the proliferative capacity. By contrast, addition of MHC-II^pos^ neutrophils from the responder animal to PBMCs (ratio 1:1) strongly suppressed T-cells as the proliferation was completely inhibited for 3 animals, and only 16% of proliferation remained for 1 animal, as compared to control wells (Fig. 6D). In one experiment we could separate MHC-II^pos^ neutrophils from the proliferating PBMCs in a transwell device and observed that suppression was abolished, indicating that the suppressive activity of MHC-II^pos^ neutrophils was contact-dependent as in the mouse (Fig 6D). Therefore, both in the mouse and the bovine, a subset of MHC-II^pos^ neutrophils can be distinguished from classical neutrophils as displaying suppressive activity on T-cells at steady state.

## DISCUSSION

We reported here that a population of MHC-II^pos^ neutrophils was present in the BM reservoir, circulated in blood and was enriched in lymphoid organs, in the apparently healthy bovine and mouse. Both MHC-II^pos^ and MHC-II^neg^ neutrophils displayed the polylobed nucleus, and were undistinguishable by this gold standard of neutrophils characterization[14]. Similar to classical MHC-II^neg^ neutrophils, the MHC-II^pos^ subset displayed important functions of neutrophils such as ROS production, phagocytosis and bacterial killing. However, unlike classical MHC-II^neg^ neutrophils, the MHC-II^pos^ neutrophils were able to suppress T-cells, a function that is reported here for the first time for bovine neutrophils. Heterogeneity or plasticity of neutrophils has largely emerged in the literature in humans or mice and new models of neutrophil differential development are proposed[15]. Here, mouse and bovine MHC-II^pos^ suppressive neutrophils were detected in the BM reservoir. They were the only ones to exert suppressive activity on T cells. They displayed a clearly distinct transcriptomic profile as compared to MHC-II^neg^ neutrophils. They were also highly enriched in lymphoid organs. Thus, they may represent a distinct subset, produced in the BM for rapid mobilization and regulation of T cells. On the other hand, CD11b upregulation that was observed on these neutrophils could also sign an activated state[29]. Recently, CD11b^hi^ primed neutrophils were also reported to circulate in blood in healthy mice and humans to quickly respond to danger[30]. Neutrophils from healthy humans could be induced *in vitro* to exert ROS, CD11b and contact-dependent suppressive activity on T cells upon activation with specific stimuli[31]. We detected MHC-II^pos^ neutrophils in the BM and, at least in the mouse, they were able to suppress T-cell proliferation, suggesting that they were already present as regulatory cells in the reservoir. Moreover, the regulatory neutrophils from the two species did not undergo higher apoptosis or death as compared to their classical neutrophils counterparts upon stimulation with LPS, indicating they were not “older” or hyperactivated[32]. In addition, MHC-II expression by MHC-II^neg^ neutrophils could not be induced *in vitro* by incubation with LPS (data not shown). Whether heterogenous neutrophils are released from the bone-marrow as distinct subsets under steady state conditions or correspond to activation or polarization states *in situ* remains an open debate[33] but what we observed here is that MHC-II^pos^ neutrophils able to regulate T cells display a specific neutrophil “phenotype”[34].

Suppressive neutrophils that may be included within a broader category of cells termed “Myeloid-Derived Suppressor Cells (MDSCs)”[35] accumulate mostly under pathological conditions. Two main branches of MDSCs have been described: the monocytic MDSCs and the granulocytic MDSCs (G-MDSCs)[35]. In cancer, G-MDSCs are associated with tumor progression and escape to the immune surveillance[36]. In chronic infections such as tuberculosis[37; 38] or AIDS[39], or acute inflammatory syndrome such as sepsis[40] MDSCs are targeted for new host-directed therapies. However, MDSCs, including regulatory neutrophils, are also beneficial to the host under some circumstances. Because they sustain the generation of regulatory T-cells[41] they may help avoiding graft rejection[42]. They can positively regulate autoimmune disorders[43] or protect the lung from destructive inflammation during *Pseudomonas aeruginosa* or *Klebsiella pneumoniae* infections[44]. They accumulate during pregnancy where they are important regulators of fetal-maternal tolerance[45] and in neonates where they could prevent overwhelming inflammation following microbial colonization[46]. Recently, Aarts and colleagues demonstrated that neutrophils from healthy humans could become T-cell suppressive under specific stimulation[31]. Along the same line, Fine *et al* recently described that neutrophils circulate in healthy donors under several states included as “primed” cells that are first recruited to the injury site to combat infection[30]. In both these studies, high levels of CD11b signed the phenotype of these neutrophils, alike the MHC-II^pos^ regulatory neutrophil subset that we describe here. In cattle, there is no description to date of suppressive neutrophils or G-MDSCs. In a recent report, Li and colleagues[47] identified a subset of MHC-II^pos^ neutrophils that accumulated in spleen during *Ostertagia ostertagi* parasitic infection in cattle and produced the suppressive cytokine IL-10. This subset was the master regulator of immune suppression in parasitized animals. We did not observe *il10* transcription in any neutrophil subset nor IL-10 production in the supernatant of neutrophil-T-cells cocultures in our study (date not shown). Whether MHC-II^pos^ regulatory neutrophils that we identified are related to the subset identified by Li et al[47] remains an open question.

Regulatory neutrophils, or G-MDSCs, block T-cell proliferation through different mechanisms[14] including ROS production[48]. We observed in both species that direct contact between MHC-II^pos^ neutrophils and T-cells was necessary to induce suppression, as was also reported for G-MDSCs. Formation of a synapse is believed to greatly enhance suppression especially when the delivery of short-lived molecules such as H_2_O_2_ is instrumental[35]. Interestingly, we observed here a significantly higher production of total ROS upon chemical stimulation by MHC-II^pos^ neutrophils as compared to their MHC-II^neg^ counterparts which did not translate into higher killing of *E. coli*. Pliyev and colleagues demonstrated that NADPH dependent ROS were required for the expression of MHC-II in human neutrophils activated by GM-CSF and IFNγ[49]. Whether the higher ROS production recorded in our study is linked to the suppression mechanism remains to be investigated. In both the mouse and the bovine (data not shown), MHC-II was also involved in the mechanism, as blocking this molecule partially relieved the suppression of T-cell proliferation, emphasizing the need for synapse formation between the T-cell and the regulatory neutrophil. In the mouse, we observed that CD11b was also involved, as reported for human neutrophils acquiring suppressive activity upon *in vitro* stimulation[31]. Another feature of mouse regulatory neutrophil was surface expression and up-regulated gene transcription of *PD-L1*. This molecule is an important immune check-point and a favorite target for cancer treatment. MDSCs express high levels of PD-L1[50] which is involved in T-cell suppression. We could not investigate PD-L1 expression by cattle MHC-II^pos^ regulatory neutrophils due to a lack of reagents. Of note, CD14^pos^ monocytes expressing PD-L1 accumulate in blood of cattle infected by *Mycoplasma bovis*[51], a strongly immunosuppressive pathogen causing antibiotic resistant mastitis ant other diseases. PD-L1 is now a target for host-directed therapies of cattle infected by this pathogen[52].

To conclude, comparative analysis of mouse and bovine species allowed us to characterize a new subset of regulatory neutrophils that are able to suppress T cells. In the near future, we will investigate how these cells behave during clinical conditions in cattle such as mastitis, which remains one of the most important issue in dairy farming. We believe that such studies are of utmost importance to better understand the physiopathology of this disease, especially during chronic infections that remain difficult to treat. Our findings could lead to the discovery of new biomarkers and development of innovative host-directed therapies[52] targeting regulatory neutrophils for more effective clearance of pathogens and better control of mammary gland inflammation and damage.

## ACKNOWLEDGMENTS

Marion Rambault conducts her doctoral project under a CIFRE agreement (Industrial agreement of learning/training by research, CIFRE N° 2019/0776) signed with Institut de l’Elevage IDELE. We are grateful to Théodore Vinais for his early contribution during his internship to the characterization of the two neutrophils subsets in cattle.

Corinne Beauge and her team (PFIE, INRAE, Nouzilly) are gratefully acknowledged for mice care. We warmly thank Eric Briant and his team (UE-PAO, INRAE, Nouzilly) for bovine blood sampling and animal care. We thank the staff of the Abattoir du Perche Vendômois for valuable access to and assistance for bovine post-mortem sampling. Bovine CD11b hybridoma was a kind gift from Dr. Dirk Werling (Royal Veterinary College of London).

This work was supported by a grant from the French Agence Nationale de la Recherche under the Carnot Program France Future Elevage (BoNeutro), the EGER program of APIS-GENE (MASTICELLS), and the Région Centre Val de Loire grant N°32000584 “inflammation et infection”) and the EUROFERI project (FEDER-FSE Centre Val de Loire 2014-2020, N° EX 010233).

## AUTHOR CONTRIBUTIONS

MR designed and did most of the experiments for cattle neutrophils and EDD for mouse neutrophils. They analyzed data and prepared all manuscript figures. YLV realized flow cytometry analysis and sorting of mouse and cattle neutrophils. PC helped with the Mixed Leukocyte Reaction in cattle. FG helped with bacterial experiments. PG helped with transcriptomic analysis, critically analyzed the data and revised the manuscript. PR brought valuable expertise on cattle neutrophils purification methods, critically analyzed the data and revised the manuscript. NW and AR obtained grants, supervised all aspects of the work, critically analyzed the data and wrote the manuscript. AR also contributed to important experiments in cattle. All authors read and approved the manuscript for publication.

## Supplementary table and figures legends

**Supplementary Table S1.** Antibodies used in the study. The specificity, origin, clone number and commercial provenance of all antibodies used in the study are listed.

|  |  | Target | Reactivity | Clone | Supplier | Isotype | Dye |
| --- | --- | --- | --- | --- | --- | --- | --- |
| Mouse |  | CD11b | mouse | M1/70 | BD biosciences | Rat IgG<sub>2b</sub>, κ | V450 |
|  |  | Ly6G | mouse | 1A8 | BD biosciences | Rat IgG<sub>2a</sub>, κ | PE |
|  |  | Ly6C | mouse | AL-21 | BD biosciences | Rat IgM, κ | APC |
|  |  | MHC-II | mouse | 2G9 | BD biosciences | Rat IgG<sub>2a</sub>, κ | FITC |
|  |  | CD14 | mouse | rmC5-3 | BD biosciences | Rat IgG1, κ | BV711 |
|  |  | CD62-L | mouse | MEL-14 | BD biosciences | Rat IgG<sub>2a</sub>, κ | BV711 |
|  |  | CD44 | mouse | IM7 | BD biosciences | Rat IgG<sub>2b</sub>, κ | BV711 |
|  |  | CD274 | mouse | MIH5 | BD biosciences | Rat IgG<sub>2a</sub>, I | BV711 |
| Bovine | Primary Ab | Granulocytes | bovine | CH138A | KingFisher Biotech | Mouse IgM | - |
|  |  | MHC-II | feline | CAT82A | KingFisher Biotech | Mouse IgG<sub>1</sub> | - |
|  |  | CD14 | bovine | CC-G33 | Bio-Rad | Mouse IgG<sub>1</sub> | - |
|  |  | CD62L | bovine | FMC46 | OriGene | Mouse IgG<sub>2b</sub> | - |
|  |  | CD11b (hybridoma) | bovine | AFRC IAH CC104 | See acknowledgment | Mouse IgG<sub>2b</sub> | - |
|  | Secondary Ab | IgM | mouse | - | Invitrogen | Goat IgG | Alexa Fluor 647 |
|  |  | IgG<sub>1</sub> | mouse | - | Invitrogen | Goat IgG | Alexa Fluor 488 |
|  |  | IgG<sub>2b</sub> | mouse | RMG2b-1 | BioLegend | Rat IgG<sub>1</sub>, κ | Phycoerythrin (PE) |
|  |  | Viability dye | - | - | eBiosciences | - | eFluor780 |
|  |  | Annexin V | Mouse, bovine | - | BD biosciences | - | Biotin or FITC |
|  |  | Streptavidin | - | - | BD Biosciences | - | APC-Cy7 |

**Supplementary Table S2.**
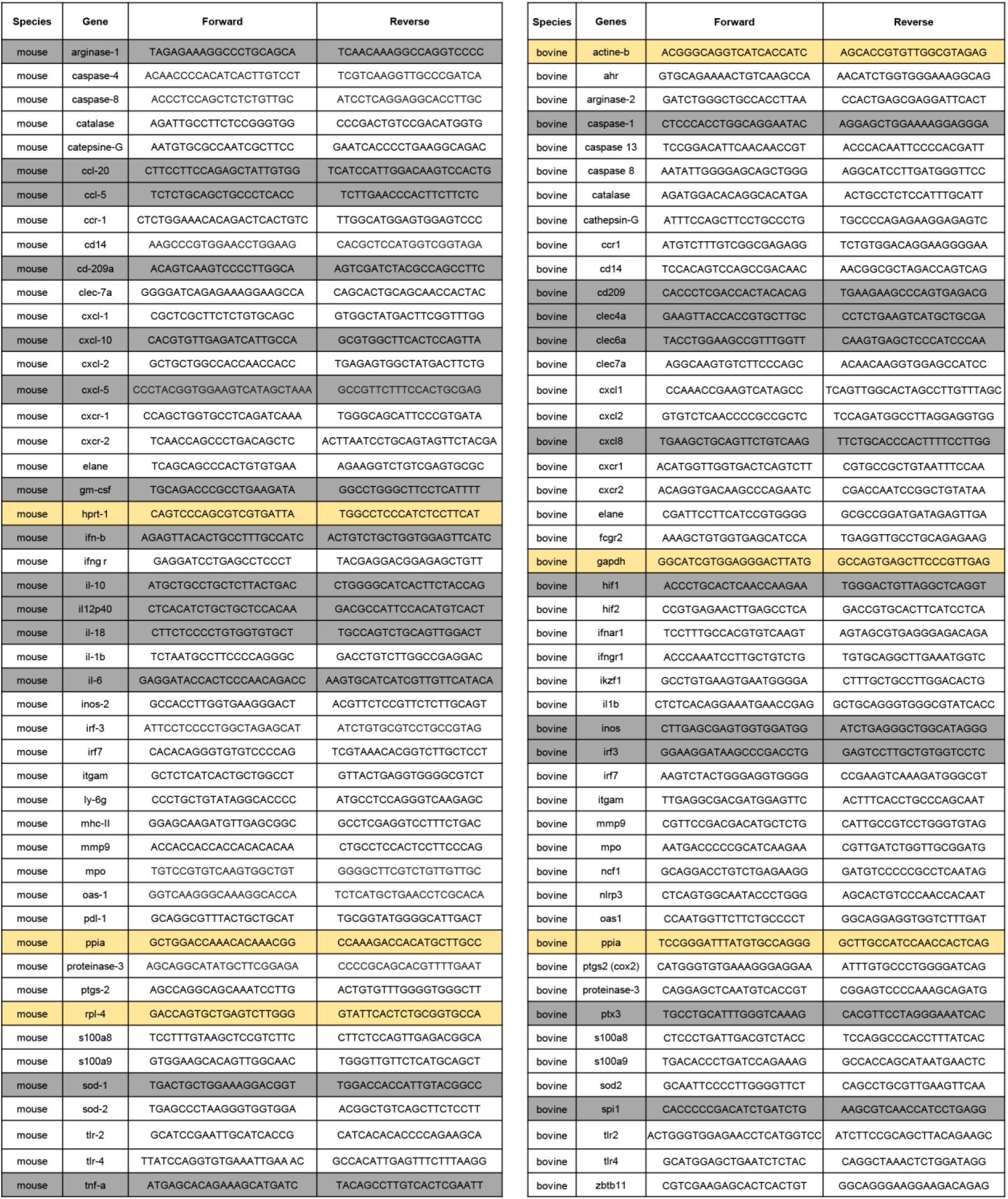
Sequences of primers used in this study. Primers were designed using Geneious software, in intron-spanning regions when possible. The annealing temperature was set at 60 and 62°C for bovine and mouse samples respectively. Housekeeping genes used as the reference to calculate ΔCT for each species are indicated in the yellow boxes and weakly expressed genes that were removed from the Principal Component Analysis presented in Fig 4 are indicated in the grey boxes.

**Supplementary Figure S1.**
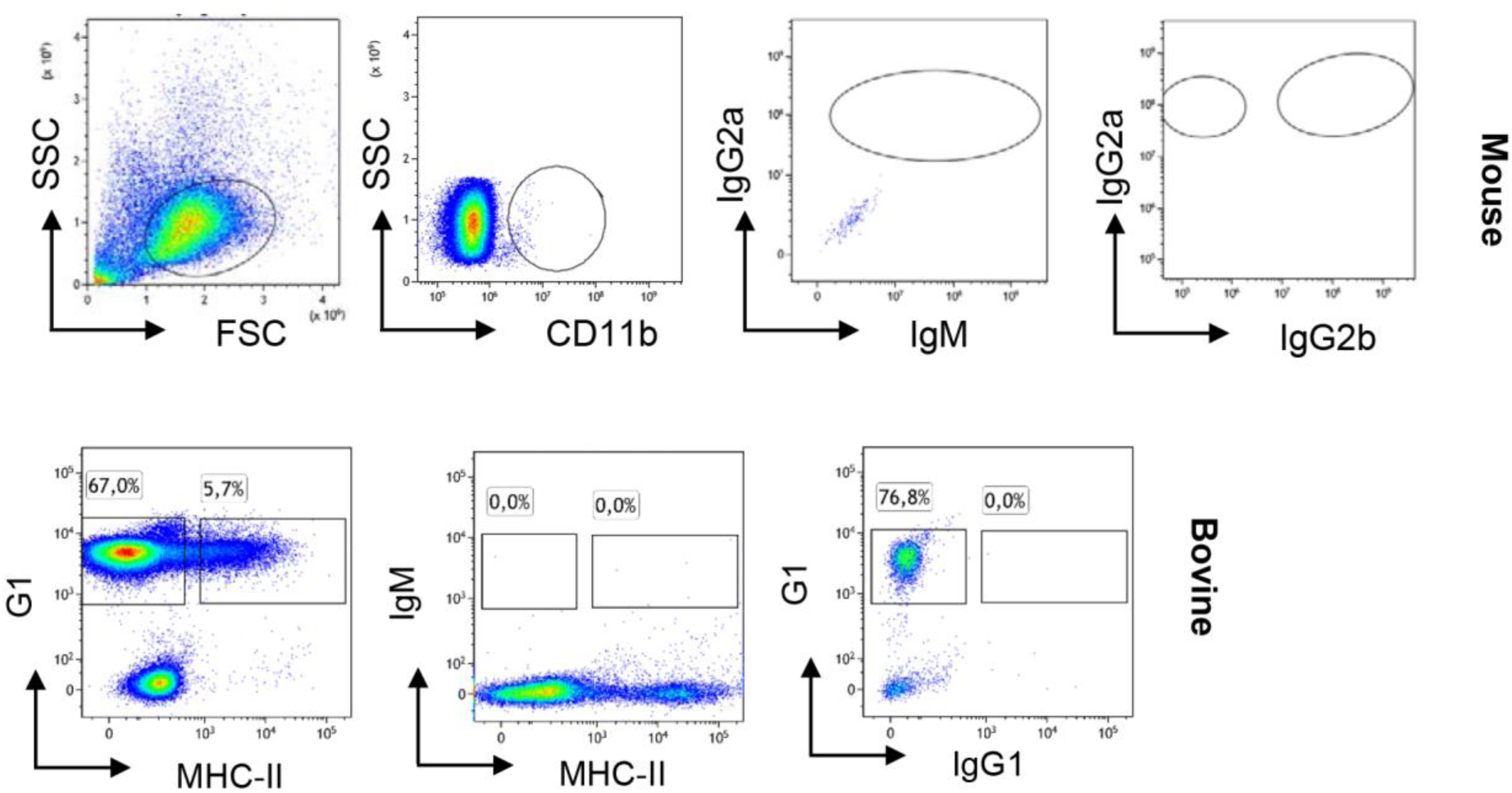
Isotype controls for neutrophil diversity analysis in mouse bone marrow and cattle blood. Mouse and bovine neutrophils were labelled as described in Fig. 2. Similar procedures wzere set up with isotype controls for all antibodies in each experiment to correctly set the analysis and sorting gates. Dot plots from one representative animal are depicted (3 independent experiments, n=4 mice, n=6 for bovine).

**Supplementary Figure S2.**
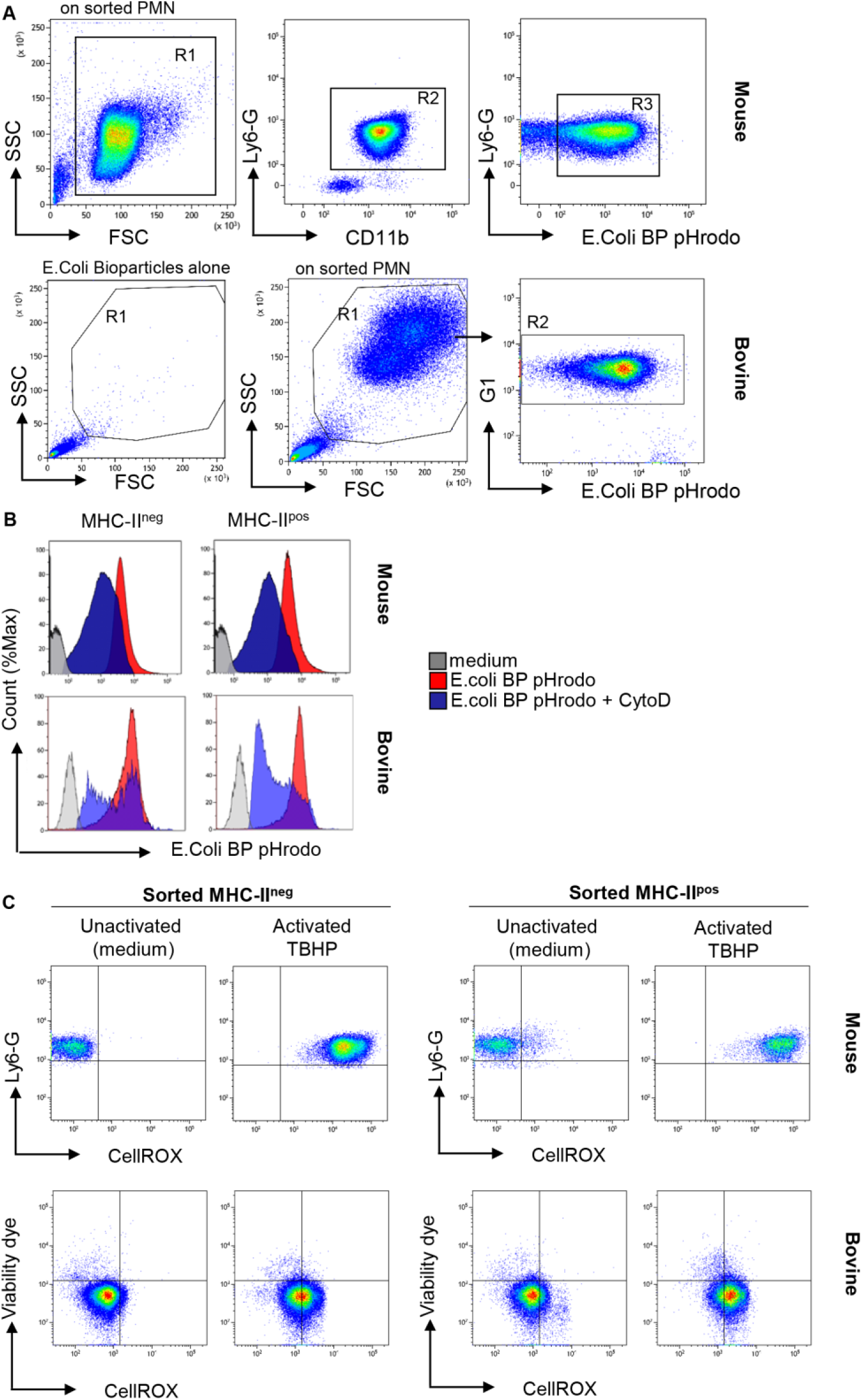
Gating strategy for analysis of phagocytosis and ROS production by neutrophils. Neutrophils were labelled and sorted as described in Fig. 2 and S1. **(A)** After purification by cell sorting from the BM (mouse) or blood (bovine) phagocytosis by MHC-II^pos^ or MHC-II^neg^ neutrophils was assessed using pHrodo E.coli bioparticles with or without previous treatment with cytochalasin D. Dot plots from one representative animal are depicted (3 independent experiments, n=3 pool of 10 mice, n=3 for bovine). **(B)** Oxidative stress was measured in MHC-II^pos^ and MHC-II^neg^ sorted neutrophils using the CellROX Orange probe that reacts with all ROS species. Cells were activated with TBHP or incubated with medium alone and levels of ROS were measured by flow cytometry among the live cells (unstained with eFluor780 viability dye). Dot plots from one representative animal are depicted (3 and 4 independent experiments for mice and cattle respectively, n=3 pool of 10 mice, n=5 for bovine).

**Supplementary Figure S3.**
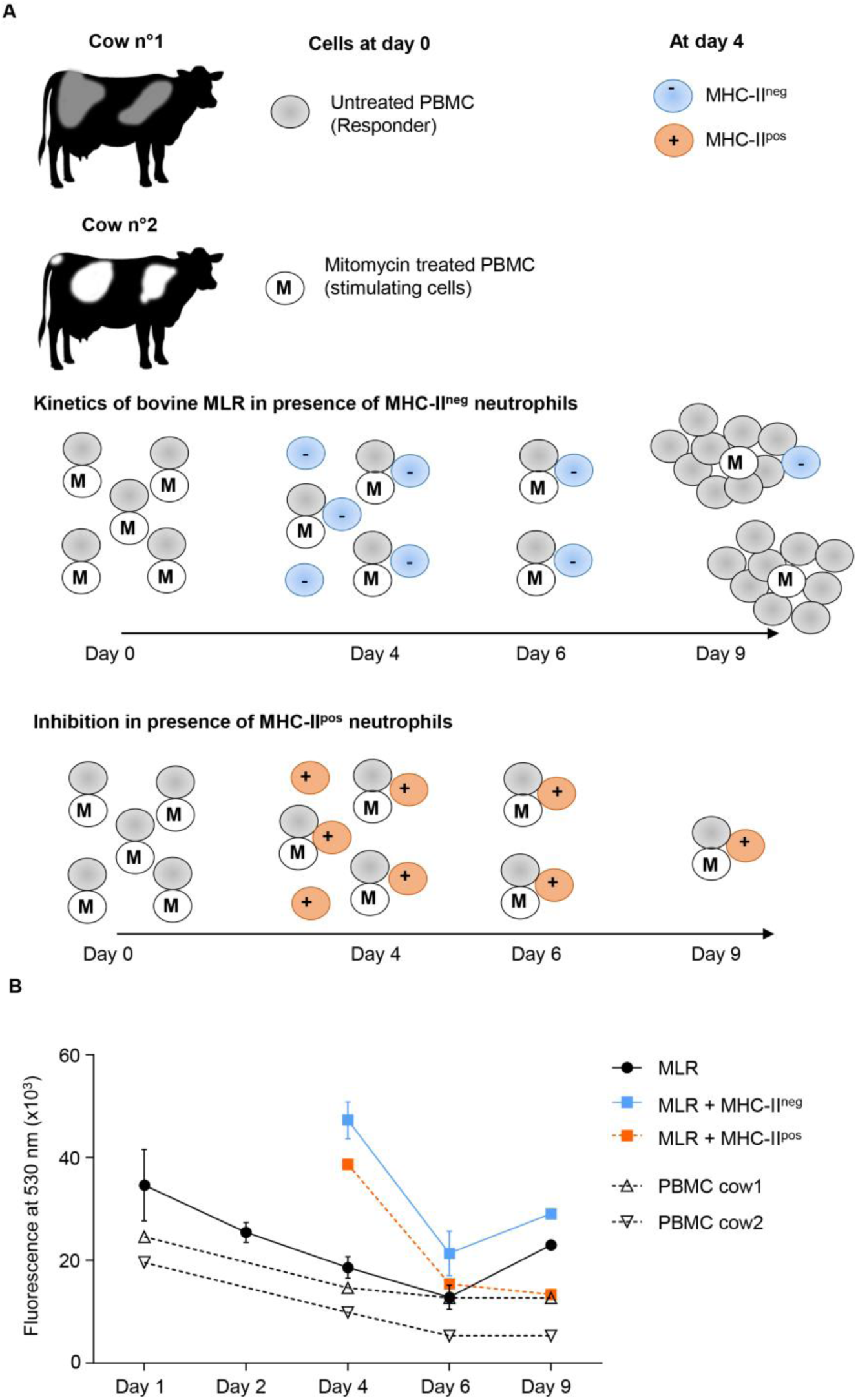
Kinetics of bovine Mixed Leukocyte Reaction and analysis of neutrophil suppressive activity. **(A)** PBMCs from the responder animal were isolated and left untreated, while PBMCs from the stimulating animal were incubated with mitomycin C to block their proliferation. PBMCs from the two cows were incubated at ratio of 1:1. Sorted MHC-II^pos^ or MHC-II^neg^ neutrophils from the responder animal were added to the reaction at day 4. **(B)** DNA was quantified at different time points with CyQUANT Cell Proliferation Assay tests according to manufacturer’s instruction and fluorescence was read at 530nm. DNA extracted from PBMCs cultivated separately decreased along the assay indicated the absence of proliferation (dotted lines). In the MLR reaction, while DNA content declined between day 1 and 6, PBMCs proliferation could be measured between day 6 and 9 (black). The effect of adding sorted MHC-II^neg^ neutrophils (blue) or MHC-II^pos^ neutrophils (orange) to the proliferating cells could then be measured. One representative experiment is shown and data represent the mean ± SEM of technical triplicates. Four independent experiments were conducted with different pairs of cows.

